# Abundant contribution of short tandem repeats to gene expression variation in humans

**DOI:** 10.1101/017459

**Authors:** Melissa Gymrek, Thomas Willems, Haoyang Zeng, Barak Markus, Mark J. Daly, Alkes L. Price, Jonathan Pritchard, Yaniv Erlich

## Abstract

Expression quantitative trait loci (eQTLs) are a key tool to dissect cellular processes mediating complex diseases. However, little is known about the role of repetitive elements as eQTLs. We report a genome-wide survey of the contribution of Short Tandem Repeats (STRs), one of the most polymorphic and abundant repeat classes, to gene expression in humans. Our survey identified 2,060 significant expression STRs (eSTRs). These eSTRs were replicable in orthogonal populations and expression assays. We used variance partitioning to disentangle the contribution of eSTRs from linked SNPs and indels and found that eSTRs contribute 10%-15% of the *cis*-heritability mediated by all common variants. Functional genomic analyses showed that eSTRs are enriched in conserved regions, co-localize with regulatory elements, and are predicted to modulate histone modifications. Our results show that eSTRs provide a novel set of regulatory variants and highlight the contribution of repeats to the genetic architecture of quantitative human traits.

## Introduction

In recent years, there has been tremendous progress in identifying genetic variants that affect expression of nearby genes, termed *cis* expression quantitative trait loci (*cis*-eQTLs). Multiple studies have shown that disease-associated variants often overlap *cis*-eQTLs in the affected tissue^1,2^. These observations suggest that understanding the genetic architecture of the transcriptome may provide insights into the cellular-level mediators underlying complex traits^3-5^. So far, eQTL-mapping studies have mainly focused on SNPs and to a lesser extent on bi-allelic indels and CNVs as determinants of gene expression^6-10^. However, these variants do not account for all of the heritability of gene expression attributable to *cis*-regulatory elements as measured by twin studies, leaving on average about 20-30% unexplained^7,11^. It has been speculated that such heritability gaps could indicate the involvement of repetitive elements that are not well tagged by common SNPs^12,13^.

To augment the repertoire of eQTL classes, we focused on Short Tandem Repeats (STRs), one of the most polymorphic and abundant type of repetitive elements in the human genome^14,15^. These loci consist of periodic DNA motifs of 2-6bp spanning a median length of around 25bp. There are about 700,000 STR loci covering almost 1% of the human genome. Their repetitive structure induces DNA-polymerase slippage events that add or delete repeat units, creating mutation rates that are orders of magnitude higher than those of most other variant types^14,16^. Over 40 Mendelian disorders, such as Huntington’s Disease, are attributed to STR mutations, most of which are caused by large expansions of trinucleotide coding repeats^17^. However, trinucleotide coding STRs are only a minute fraction of all genomic STRs. The majority consist of di- and tetranucleotide motifs, which are overrepresented in promoter and regulatory regions^18^.

Multiple lines of evidence support the potential role of STRs in regulating gene expression. *In vitro* studies have shown that STR variations can modulate the binding of transcription factors^19,20^, change the distance between promoter elements^21,22^, alter splicing efficiency^23,24^, and induce irregular DNA structures that may modulate transcription^25^. Recent computational work showed that dinucleotide STRs are a hallmark of enhancer elements in *Drosophila*^26^. *In vivo* experiments have reported specific examples of STR variations that control gene expression across a wide range of taxa, including *Haemophilus influenza^27^*, *Saccharomyces cerevisiae^28^*, *Arabidopsis thaliana^29^*, and vole^30^. In humans, several dozen candidate-gene studies used reporter assay experiments to show that STR variations modulate gene expression^19,31-35^ and alternative splicing^23,36,37^. However, there has been no systematic evaluation of the contribution of STRs to gene expression in humans.

To that end, we conducted a genome-wide analysis of STRs that affect expression of nearby genes, termed expression STRs (eSTRs), in lymphoblastoid cell lines (LCLs), a central *ex-vivo* model for eQTL studies. This well-studied model permitted the integration of whole genome sequencing data, expression profiles from RNA-sequencing and arrays, and functional genomics data. We tested for association in close to 190,000 STR×gene pairs and found over 2,000 significant eSTRs. Using a multitude of statistical genetic and functional genomics analyses, we show that hundreds of these eSTRs are predicted to be functional, uncovering a new class of genetic variants that modulate gene expression.

## Results

### Initial genome-wide discovery of eSTRs

The initial genome-wide discovery of potential eSTRs relied on finding associations between STR length and expression of nearby genes. We focused on 311 European individuals whose LCL expression profiles were measured using RNA-sequencing by the gEUVADIS^8^ project and whose whole genomes were sequenced by the 1000 Genomes Project^38^. The STR genotypes were obtained in our previous study^39^ in which we created a catalog of STR variation as part of the 1000 Genomes Project using lobSTR, a specialized algorithm for profiling STR variations from high throughput sequencing data^40^. Briefly, lobSTR identifies reads with repetitive sequences that are flanked by non-repetitive segments. It then aligns the non-repetitive regions to the genome using the STR motif to narrow the search, thereby overcoming the gapped alignment problem and conferring alignment specificity. Finally, lobSTR aggregates aligned reads and employs a model of STR-specific sequencing errors to report the maximum likelihood genotype at each locus. lobSTR recovered most (r^2^=0.71) of the additive genetic variance of STR loci in the 1000 Genomes datasets based on large-scale validation using 5,000 STR genotype calls obtained by capillary electrophoresis, the gold standard for STR genotyping^39^. The majority of genotype errors were from dropout of one allele at heterozygote sites due to low sequencing coverage. We simulated the performance of STR associations using lobSTR calls compared to the capillary calls. This process showed that STR genotype errors reduce the power to detect eSTRs by 30-50% but importantly do not create spurious associations (**Supplementary Note** and **Supplementary Fig. 1**).

**Figure 1:**
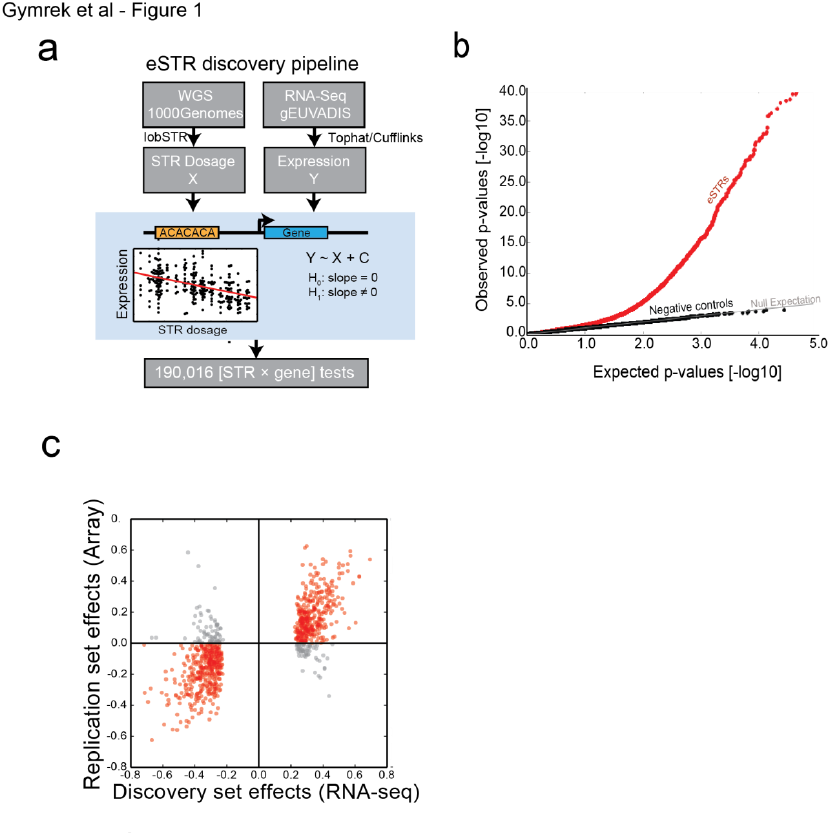
eSTR discovery and replication. **(a)** eSTR discovery pipeline. An association test using linear regression was performed between STR dosage and expression level for every STR within 100kb of a gene **(b)** Quantile-quantile plot showing results of association tests. The gray line gives the expected p-value distribution under the null hypothesis of no association. Black dots give p-values for permuted controls. Red dots give the results of the observed association tests **(c)** Comparison of eSTR effect sizes as Pearson correlations in the discovery dataset vs. the replication dataset. Red points denote eSTRs whose direction of effect was concordant in both datasets and gray points denote discordant directions.

To detect eSTR associations, we regressed gene expression on STR dosage, defined as the sum of the two STR allele lengths in each individual. We opted to use this measure based on previous findings that reported a linear trend between STR length and gene expression^19,32,34^ or disease phenotypes^41,42^. As covariates, we included sex, population structure, and other technical parameters (**Fig. 1a** and **Supplementary Methods**). We employed this process on 15,000 coding genes whose expression profiles were detected in the RNA-sequencing data. For each gene, we considered all polymorphic STR variations that passed our quality criteria (**Methods**) within 100kb of the transcription start and end sites of the gene transcripts as annotated by Ensembl^43^. On average, 13 STR loci were tested for each gene (**Supplementary Fig. 2**), yielding a total of 190,016 STR×gene tests.

**Figure 2:**
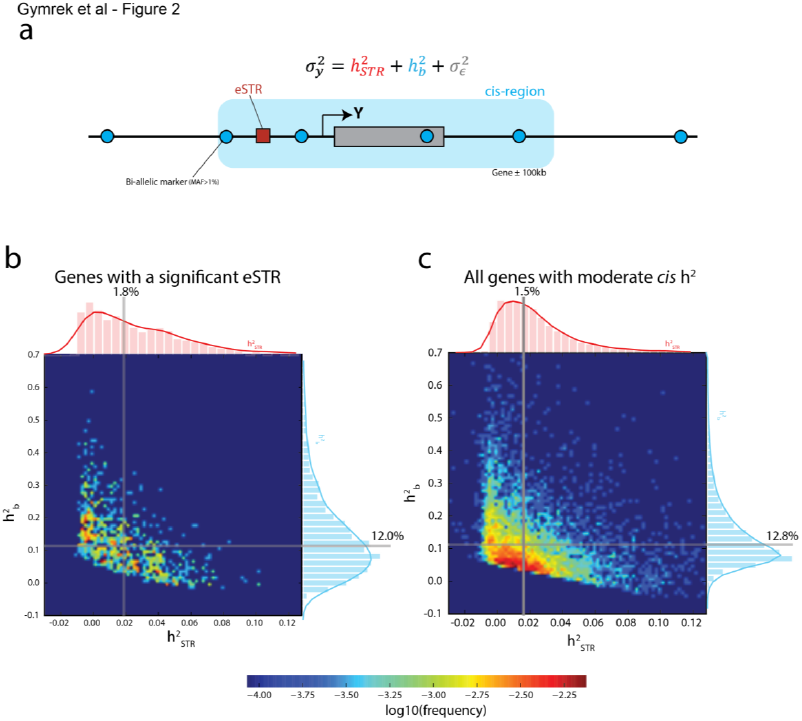
Variance partitioning using linear mixed models. **(a)** The normalized variance of the expression of gene Y was modeled as the contribution of the best eSTR and common bi-allelic markers in the *cis* region (±100kb from the gene boundaries) (**b**&**c**) Heatmaps show the joint distributions of variance explained by eSTRs and by the *cis* region. Gray lines denote the median variance explained **(b)** Variance partitioning across genes with a significant eSTR in the discovery set and **(c)** variance partitioning across genes with moderate *cis* heritability.

Our analysis identified 2,060 unique protein-coding genes with a significant eSTR (gene level FDR≤5%) (**Fig. 1b**, **Supplementary Table 1**). The majority of these were di- and tetra-nucleotide STRs (**Supplementary Tables 2**, **3**). Only 13 eSTRs fall in coding exons but eSTRs were nonetheless strongly enriched in 5’UTRs (p=1.0×10^-8^), 3’UTRs (p=1.7×10^-9^) and regions near genes (p<10^-28^) compared to all STRs analyzed (**Supplementary Table 4**). We repeated the association tests with two negative control conditions by regressing expression on (i) STR dosages permuted between samples and (ii) STR dosages from randomly chosen unlinked loci (**Fig. 1b, Supplementary Fig. 3**). Both negative controls produced uniform p-value distributions expected under the null hypothesis. This provides support for the absence of spurious associations due to inflation of the test statistic or the presence of uncorrected population structure.

**Figure 3:**
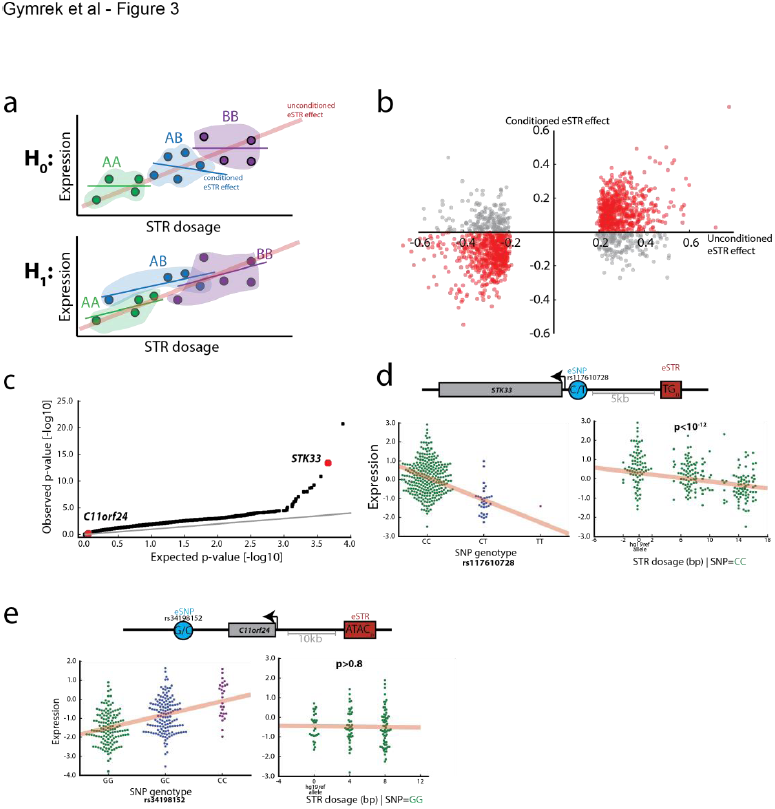
eSTR associations in the context of eSNPs. **(a)** Schematic of the eSTR effect versus the effect conditioned on the best eSNP genotype. Under the null expectation, the original association (red line) comes from mere tagging of eSNPs. Thus, the eSTR effect disappears when restricting to a group of individuals (dots) with the same eSNP genotype (colored patches). Under the alternative hypothesis, the effect is concordant between the original and conditioned associations **(b)** The original eSTR effect versus the conditioned eSTR effect. Red points denote eSTRs whose direction of effect was concordant in both datasets and gray points denote discordant directions **(c)** Quantile-quantile plot of p-values from ANOVA testing of the explanatory value of eSTRs beyond that of eSNPs (**d**) *STK33* is an example of a gene for which the eSTR (red rectangle) has a strong explanatory value beyond the best eSNP (blue circle) based on ANVOA. Indeed, when conditioning on individuals that are homozygous for the “C” eSNP allele (bottom left, green dots), the STR dosage still shows a significant effect (bottom right) **(e)** *C11orf24* is an example of a gene for which the eSTR was part of the discovery set but did not pass the ANOVA threshold. After conditioning on individuals that are homozygous for the “G” eSNP allele (bottom left, green dots), the STR effect is lost (bottom right).

The initial discovery set of eSTRs was largely reproducible in an independent set of individuals using an orthogonal expression assay technology. We obtained an additional set of over 200 individuals whose genomes were also sequenced as part of the 1000 Genomes Project and whose LCL expression profiles were measured by Illumina expression array^44^. These individuals belong to cohorts with African, Asian, European, and Mexican ancestry, enabling testing of the associations in a largely distinct set of populations. The Illumina expression array allowed testing 882 eSTRs out of the 2,060 identified above. The association signals of 734 of the 882 (83%) tested eSTRs showed the same direction of effect in both datasets (sign test p=2.7×10^-94^) and the effect sizes were strongly correlated (R=0.73, p=1.4×10^-149^) (**Fig. 1c**), despite only moderate reproducibility of expression profiles across platforms (**Supplementary Note** and **Supplementary Fig. 4**). Overall, these results show that eSTR association signals are robust and reproducible across populations and expression assay technologies.

**Figure 4:**
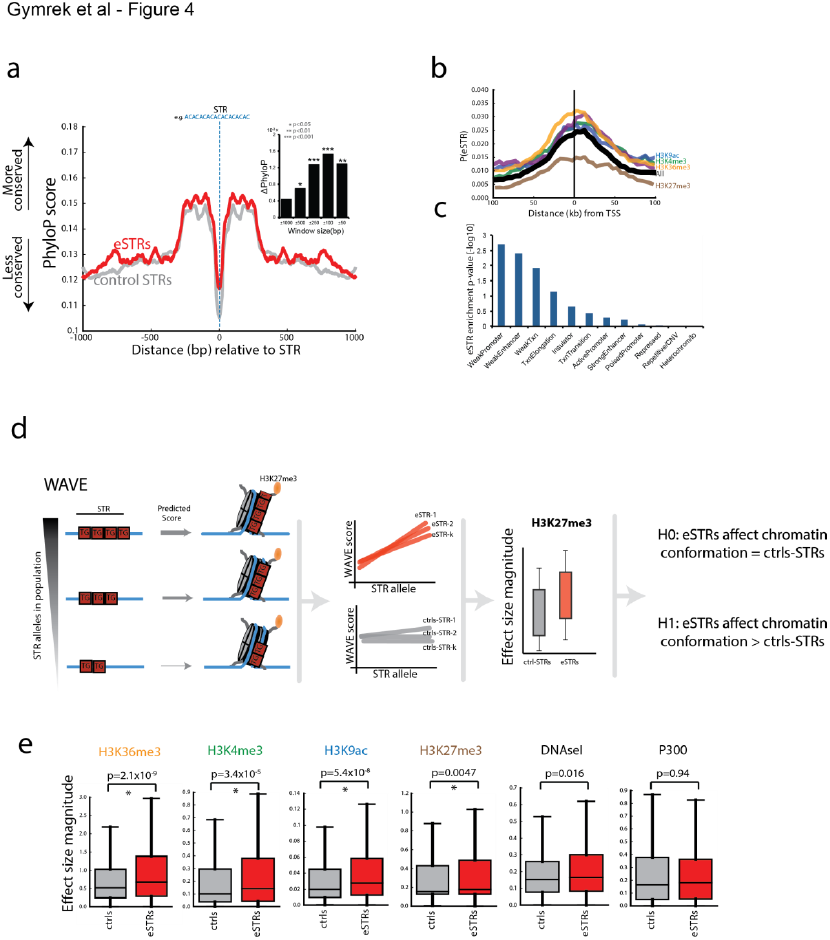
Functional analysis of eSTR loci. **(a)** Median PhyloP conservation score as a function of distance from the STR. Red: eSTR loci, gray: matched control STRs. Inset: the difference in the PhyloP conservation score between eSTRs and matched control STRs as a function of window size around the STR. **(b)** The probability that an STR scores as an eSTR in the discovery set as a function of distance from the transcription start site (TSS). eSTRs show clustering around the TSS (black line). Conditioning on the presence of a histone mark (colored lines) significantly modulated the probability of an STR to be an eSTR **(c)** The enrichment of eSTRs in different chromatin states **(d)** Schematic of the application of WAVE to predict histone modification signatures for different STR alleles. For each eSTR (red) and control STR (gray) we measured the magnitude of the slope between the STR allele and the WAVE score and then tested whether the magnitudes were significantly different between the two sets **(e)** Comparison of the distribution of slope magnitudes for eSTRs (red) and controls (gray). A “*” denotes p-values < 0.01.

### Partitioning the contribution of eSTR and nearby variants

An important question is whether eSTR association signals stem from causal STR loci or are merely due to tagging SNPs or other variants in linkage disequilibrium (LD). Previous results reported that the average STR-SNP LD is approximately half of the traditional SNP-SNP LD^39,45^, but there are known examples of STRs tagging GWAS SNPs^46^.

To address this question, we partitioned the relative contributions of eSTRs versus all common (MAF≥1%) bi-allelic SNPs, indels, and structural variants (SV) in the *cis* region using a linear mixed model (LMM) (**Fig. 2a**). Multiple studies have used this approach to measure the total contributions of common variants to the heritability of quantitative traits and to partition the contribution of different classes of variants^47,48^. Taking a similar approach, we included two types of effects for each gene: a random effect (h^2^_b_) that captures all common bi-allelic loci detected within 100kb of the gene and a fixed effect (h^2^_STR_) that captures the best STR. To test whether other causal variants on the local region could inflate the estimate of the STR contribution, we simulated gene expression with a causal SNP eQTL per gene while preserving the local haplotype structure. In this negative control scenario, the LMM correctly reported a median h^2^_STR_/h^2^_cis_≈0 across all conditions (**Supplementary Note** and **Supplementary Fig. 5**), where h^2^_cis_= h^2^_b_+h^2^_STR_. This suggests that other causal variants in LD do not inflate the estimator of the relative contribution of STRs. As the LMM is expected to downwardly bias the variance explained in the presence of genotyping errors, the reported h^2^_STR_ is likely to be conservative.

The LMM results showed that eSTRs contribute about 12% of the genetic variance attributed to common *cis* polymorphisms. For genes with a significant eSTR, the median h^2^_STR_ was 1.80%, whereas the median h^2^_b_ was 12.0% **(Fig. 2b)**, with a median ratio of h^2^_STR_/h^2^_CIS_ = 12.3% (CI_95%_ 11.1%-14.2%; n=1,928) (**Table 1**). We repeated the same analysis for genes with at least moderate (≥5%) *cis*-heritability (**Methods**) regardless of the presence of a significant eSTR in the discovery set. The motivation for this analysis was to avoid potential winner’s curse^49^ and to obtain a transcriptome-wide perspective on the role of STRs in gene expression (**Fig. 2c**). In this set of genes, eSTRs contribute about 13% (CI_95%_ 12.2%-13.4%; n=6,272) of the genetic variance attributed to *cis* common polymorphisms. The median h^2^_STR_ was 1.45% of the total expression variance, whereas the median h^2^_b_ was 9.10% (**Table 1**). Repeating the analysis while treats STRs as a random effect showed highly similar results (**Supplementary Note**, **Supplementary Table 5, Supplementary Fig. 5-6,**). Taken together, this analysis shows that STR variations explain a sizeable component of gene expression variation after controlling for all variants that are well tagged by common bi-allelic markers on in the *cis* region.

**Table 1:**
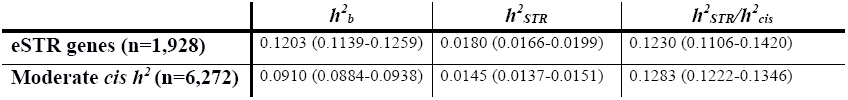
Heritability of gene expression explained by STRS vs. common bi-allelic variants. Values show the median and 95% confidence interval of the median across all eSTR-containing genes and genes with moderate *cis* heritability (=5%). *h*^2^_*b*_ denotes the variance explained by all common *cis* bi-allelic variants,*h*^2^_*STR*_ denotes the variance explained by the best STR for each gene, and *h*^2^_*cis*_= *h*^2^_*STR*_ + *h*^2^_*b*_.

### The effect of eSTRs in the context of individual SNP eQTLs

To further assess the contribution of eSTRs in the context of other variants, we also inspected the relationship between eSTRs and individual cis-SNP eQTLs (eSNPs). We performed a traditional eQTL analysis with the whole genome sequencing data of 311 individuals that were part of the discovery set to identify common eSNPs [minor allele frequency (MAF) ≥5%] within 100kb of the gene. This process identified 4,290 genes with an eSNP (gene-level FDR≤5%). We then re-analyzed the eSTR association signals while conditioning on the genotype of the most significant eSNP (**Fig. 3a**). For each eSTR, we ascertained the subset of individuals that were homozygous for the major allele of the best eSNP in the region. If the eSTR simply tags this eSNP, its conditioned effect should be randomly distributed compared to the unconditioned effect. Alternatively, if the eSTR is causal, the direction of the conditioned effect should match the original effect. We conducted this analysis for eSTR loci with at least 25 individuals homozygous for the best eSNP and for which these individuals had at least two unique STR genotypes (1,856 loci). After conditioning on the best eSNP, the direction of effect for 1,395 loci (75%) was identical to that in the original analysis (sign test p<4.2×10^-109^) and the effect sizes were significantly correlated (R=0.52; p=3.2×10^-130^) (**Fig. 3b**). This further supports the additional role of eSTRs beyond traditional cis-eQTLs.

We also found that hundreds of eSTRs in the discovery set provide additional explanatory value for gene expression beyond the best eSNP. In 23% of genes, the eSTR significantly improved the explained variance of gene expression over considering only the best eSNP according to an ANOVA model comparison (FDR<5%) (**Fig. 3c–e, Methods**). Combined with the 183 genes with an eSTR but no significant eSNP, these results show that at least 30% of the eSTRs identified by our initial scan cannot be explained by mere tagging of the best eSNP. Given the reduced quality of STR compared to SNP genotypes, this analysis is likely to underestimate the true contribution of STRs. Nonetheless, our results show concrete examples for hundreds of associations in which the eSTR increases the variance explained by the best eSNP.

### Functional Genomics Supports the Causal Role of eSTRs

To provide further evidence of their causal role, we analyzed eSTRs in the context of functional genomics data. First, we assessed the potential functionality of STR regions by measuring signatures of purifying selection, since previous findings have reported that putatively causal eSNPs are slightly enriched in conserved regions^50^. We inspected the sequence conservation^51^ across 46 vertebrates in the sequence upstream and downstream of the eSTRs in our discovery dataset (**Fig. 4a**). To tune the null expectation, we matched each tested eSTR to a random STR that did not reach significance in the association analysis but had a similar distance to the nearest transcription start site (TSS). The average conservation level of a ±500bp window around eSTRs was slightly but significantly higher (p<0.03) than for control STRs. Tightening the window size to shorter stretches of ±50bp showed a more significant contrast in the conservation scores of the eSTRs versus the control STRs (p<0.01) (**Fig. 4a inset**), indicating that the excess in conservation comes from the vicinity of the eSTR loci. Taken together, these results show that eSTRs discovered by our association pipeline reside in regions exposed to relatively higher purifying selection, further suggesting a functional role.

We also found that eSTRs significantly co-localize with functional elements. eSTRs show the strongest enrichment closest to transcription start sites (**Fig. 4b**) and to a lesser extent near transcription end sites (**Supplementary Fig. 7**), similar to patterns previously observed for eSNPs^50^. We then inspected the co-localization of eSTRs with histone modifications as annotated by the Encode Consortium^6^ in LCLs. eSTRs were strongly enriched in peaks of histone modifications associated with regulatory regions (H3K4me1, H3K4me2, H3K4me3, H3K27ac, H3K9ac) and transcribed regions (H3K36me3), and highly depleted in repressed regions (H3K27me3) (**Fig. 4b**). These results match previous patterns of enrichments found for putatively causal eSNPs^50^. To test the significance of these signals, we constructed a null distribution for each histone modification by measuring the co-localization of eSTRs with randomly shifted histone peaks similar to the fine-mapping procedure of Trynka *et al.*^52^. This null distribution controls for co-occurrence of eSTRs and histone peaks due to their proximity to other causal variants. We found eSTR/histone co-localizations were significant (weakest p-value<0.01) after the peak shifting procedure, suggesting that these results stem from the eSTRs themselves. We also performed a peak-shifting analysis using ChromHMM annotations^53^ (**Fig. 4c**). The two strongest enrichments for eSTRs were weak-promoters (p<0.002) and weak-enhancers (p<0.004). Again, this analysis shows overlap of eSTRs with elements that are predicted to regulate gene expression.

Finally, we found that eSTR variations are likely to modulate the occupancy of certain histone marks. For each eSTR, we created a series of DNA sequences reflecting the STR alleles observed among individuals in our dataset (**Fig. 4d**). We used these sequences as an input to the WAVE (Whole-genome regulAtory Variants Evaluation) model^54^, which predicts ChIP-sequencing experiments directly from genomic sequences (**Methods**). The output of WAVE showed the predicted effect of STR variations on the occupancy of chromatin marks. We then compared the distribution of the magnitude of effect sizes between eSTRs and a randomly chosen control set of STRs. eSTRs had significantly greater effects than control STRs on the predicted occupancy of all tested histone marks (p_H3K4me3_=3.4×10^-5^, p_H3K9ac_ =5.4×10^-8^, p_H3K36me3_=2.1×10^-9^, p_H3K27me3_=0.0047; Mann-Whitney rank test) (**Fig. 4e)**. We also discovered a marginally significant effect on DNAseI (p=0.016) but not on P300 (p=0.94). Importantly, since the input material for this analysis is solely STR variations that are independent of any linked variants, these results provide an orthogonal piece of evidence of the functionality of eSTRs and suggest modulating histone marks as a potential mechanism.

## Discussion

Our study conducted the first genome-wide characterization of the effect of STR variation on gene expression and identified over 2,000 potential eSTRs. Further statistical analysis showed that eSTRs contribute on average about 10-15% of the *cis-*heritability of gene expression attributed to common (MAF≥1%) polymorphisms and that at least a third of these eSTRs improve the explained heritability beyond the strongest SNP-eQTLs. Functional genomics analyses provide further support for the predicted causal role of eSTRs.

We hypothesize that there are more eSTRs to find in the genome. Variance partitioning across all moderately heritable genes showed that STRs that did not reach significance still account for a sizeable component of gene expression variance. Our analysis also had several technical limitations. First, STRs show higher rates of genotype errors than SNPs, which limited our power to detect eSTRs and likely downwardly biased their estimated contribution in the LMM and ANOVA. In addition, about 10% of STR loci in the genome could not be analyzed because they are too long to be spanned by current sequencing read lengths^39^. Second, based on previous findings in humans^19,32,34^, our association tests focused on a linear relationship between STR length and gene expression. However, experimental work in yeast reported that certain loci exhibit non-linear relationships between STR lengths and expression^28^, which are unlikely to be captured in our current analysis. Finally, our association pipeline takes into account only the length polymorphisms of STRs and cannot distinguish the effect of sequence variations inside STR alleles with identical lengths (dubbed homoplastic alleles^55^). Addressing these technical complexities would probably require phased STR haplotypes and longer sequence reads that are currently beyond reach for large sample sizes. We envision that recent advancements in sequencing technologies^56^ will further expand the catalog of eSTRs.

Previous reports have highlighted the overlap between eQTLs and GWAS hits of human diseases^7,8,11^, suggesting a similar role for STRs. These loci, as well as other repetitive elements, have been largely overlooked by GWAS studies due to the technical complexities in genotyping them across a large number of samples. Analyzing the contribution of these loci in complex disease studies will require the availability of large scale whole genome sequencing data or the development of reliable imputation methods from genotyping arrays. Regardless of the technical method, our results suggest that such efforts might reveal exciting biology beyond that observed through the prism of traditional point mutations.

## Methods

### Genotype Datasets

lobSTR genotypes were generated for the phase 1 individuals from the 1000 Genomes Project as described in^39^. Variants from the 1000 Genomes Project phase 1 release were downloaded in VCF format from the project website. HapMap genotypes were used to correct association tests for population structure. Genotypes for 1.3 million SNPs were downloaded for draft release 3 from the HapMap Consortium webpage. SNPs were converted to hg19 coordinates using the liftOver tool and filtered using Plink^57^ to contain only the individuals for which both expression array data and STR calls were available. Throughout this manuscript, all coordinates and genomic data are referenced according to hg19.

### Expression Datasets

RNA-sequencing datasets from 311 HapMap lymphoblastoid cell lines for which STR and SNP genotypes were also available were obtained from the gEUVADIS Consortium. Raw FASTQ files containing paired end 100bp Illumina reads were downloaded from the EBI website. The hg19 Ensembl transcriptome annotation was downloaded as a GTF file from the UCSC Genome Browser^58,59^ ensGene table. The RNA-sequencing reads were mapped to the Ensembl transcriptome using Tophat v2.0.7^60^ with default parameters. Gene expression levels were quantified using Cufflinks v2.0.2^61^ with default parameters and supplied with the GTF file for the Ensembl reference version 71. Genes with median FPKM of 0 were removed, leaving 23,803 genes. We restricted analysis to protein coding genes, giving 15,304 unique Ensembl genes. Expression values were quantile-normalized to a standard normal distribution for each gene.

The replication set consisted of Illumina Human-6 v2 Expression BeadChip data from 730 HapMap lymphoblastoid cell lines from the EBI website. These datasets contain two replicates each for 730 unrelated individuals from 8 HapMap populations (YRI, CEU, CHB, JPT, GIH, MEX, MKK, LWK) and were generated as described by Stranger *et al.*^62^. Background corrected and summarized probeset intensities (by Illumina software) contained values for 7,655 probes. Additionally, probes containing common SNPs were removed^63^. Only probes with a one-to-one correspondence with Ensembl gene identifiers were retained. We removed probes with low concordance across replicates (Spearman correlation ≤ 0.5). In total we obtained 5,388 probes for downstream analysis.

Each probe was quantile-normalized to a standard normal distribution across all individuals separately for each replicate and then averaged across replicates. These values were quantile-normalized to a standard normal distribution for each probe.

### eQTL association testing

Expression values were adjusted for individual sex, individual membership, gene expression heterogeneity, and population structure (**Supplementary Methods**). Adjusted expression values were used as input to the eSTR analysis. To restrict to STR loci with high quality calls, we filtered the callset to contain only loci where at least 50 of the 311 samples had a genotype call. To avoid outlier genotypes that could skew the association analysis, we removed any genotypes seen less than three times. If only a single genotype was seen more than three times, the locus was discarded. To increase our power, we further restricted analysis to the most polymorphic loci with heterozygosity of at least 0.3. This left 80,980 STRs within 100kb of a gene expressed in our LCL dataset.

A linear model was used to test for association between normalized STR dosage and expression for each STR within 100kb of a gene (**Supplementary Methods**). Dosage was defined as the sum of the deviations of the STR allele lengths from the hg19 reference. For example, if the hg19 reference for an STR is 20bp, and the two alleles called are 22bp and 16bp, then the dosage is equal to (22-20)+(16-20) = -4bp. Then, STR genotypes were zscore-normalized to have mean 0 and variance 1. For genes with multiple transcripts we defined the transcribed region as the maximal region spanned by the union of all transcripts. The linear model for each gene is given by:

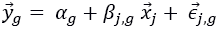

where 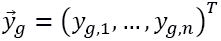 with *y*_*g,i*_ the normalized covariate-corrected expression of gene *g* in individual *i*, *n* is the number of individuals, *α*_*g*_ is the mean expression level of homozygous reference individuals, *β*_*j,g*_ is the effect of the allelic dosage of STR locus *j* on gene *g*, 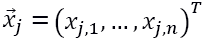 with *x*_*j*,*i*_ the normalized allelic dosage of STR locus *j* in the *i*th individual, and 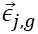 is a random vector of length *n* whose entries are drawn from 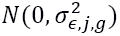 where 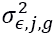 is the unexplained variance after regressing locus *j* on gene *g*. The association was performed using the OLS function from the Python statsmodels package. For each comparison, we tested *H*_0_: *β*_*j,g*_ = 0 vs. *H*_1_: *β*_*j,g*_ ≠ 0 using a standard *t*-test. We controlled for a gene-level false discovery rate (FDR) of 5% (**Supplementary Methods**).

### Partitioning heritability using linear mixed models

For each gene, we used a linear mixed model to partition heritability between the best explanatory STR and other *cis* variants. We used a model of the form:

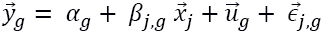

where:

- 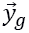, *α*_*g*_, *β*_*j,g*_, 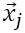, and 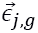 are as described above.
- 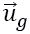 is a length *n* vector of random effects and 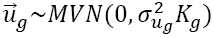 with 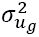 the percent of phenotypic variance explained by *cis* variants for gene *g*.
- *K*_*g*_ is a standardized *n* × *n* identity by state (IBS) relatedness matrix constructed using all common bi-allelic variants (MAF≥1%) reported by phase 1 of the 1000 Genomes Project within 100kb of gene *g*. This includes SNPs, indels, and several biallelic structural variants and is constructed as 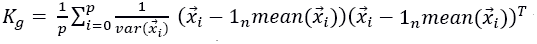 where *p* is the total number of variants considered, 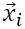 is a length *n* vector of genotypes for variant *i*, and 1_*n*_ is a length *n* vector of ones. Note the mean diagonal element of *K*_*g*_ is equal to 1.

We used the GCTA program^64^ to determine the restricted maximum likelihood estimates (REML) of *β*_*j,g*_ and 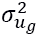. To get unbiased values of 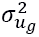, the --reml-no-constrain option was used.

We used the resulting estimates to determine the variance explained by the STR and the *cis* region. We can write the overall phenotypic variance-covariance matrix as:

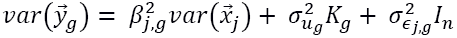

where:

- 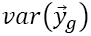 is an *n* × *n* expression variance-covariance matrix with diagonal elements equal to 1, since expression values for each gene were normalized to have mean 0 and variance 1.
- *I*_*n*_ is the *n* × *n* identity matrix.

This equation shows the relationship:

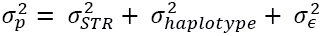

where:

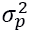 is the phenotypic variance, which is equal to 1.

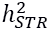 is the variance explained by the STR. This is equal to 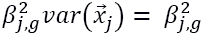 since the STR genotypes were scaled to have mean 0 and variance 1.

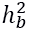 is the variance explained by bi-allelic variants in the *cis* region. This is approximately equal to 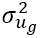 since the local IBS matrix *K*_*g*_ has a mean diagonal value of 1.

We estimated the percent of phenotypic variance explained by STRs, 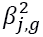, using the unbiased estimator 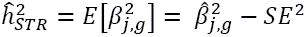, where 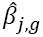 is the estimate of *β*_*j,g*_ returned by GCTA, and *SE* is the standard error on the estimate, using the fact that 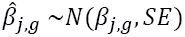 We estimated the percent of phenotypic variance explained by bi-allelic markers as 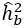. Note for this analysis the STR was treated as a fixed effect. We also reran the analysis treating the STR as a random effect, and found very little change in the results (**Supplementary Note**).

Results are reported for all eSTR-containing genes and for all genes with moderate total *cis* heritability, which we define as genes where 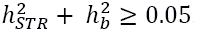. We used this approach as to our knowledge there are no published results about the *cis*-heritability of expression of individual genes in LCLs from twin studies. We used 10,000 bootstrap samples of each distribution to generate 95% confidence intervals for the medians.

### Comparing to the best eSNP

We identified SNP eQTLs using SNPs with MAF ≥ 1% as reported by phase 1 of the 1000 Genomes Project. We used an identical pipeline to our eSTR analysis to identify SNP eQTLs replacing the vector 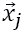 with a vector of SNP genotypes (0, 1 or 2 reference alleles) that was z-normalized to have mean 0 and variance 1. To determine whether our eSTR signal was indeed independent of the best SNP eQTL at each gene, we repeated association tests between STR dosages and expression levels while holding the genotype of the SNP with most significant association to that gene constant. For this, we determined all samples at each gene that were either homozygous reference or homozygous non-reference for the best SNP. For the SNP allele with more homozygous samples, we repeated the eSTR linear regression analysis and determined the sign and magnitude of the slope. We removed any genes for which there were less than 25 samples homozygous for the SNP genotype or for which there was no STR variation after holding the SNP constant, leaving 1,856 genes for analysis. We used a sign test to determine whether the direction of effects before and after conditioning on the best SNP are more concordant than expected by chance.

We used model comparison to determine whether eSTRs can explain additional variation in gene expression beyond that explained by the best eSNP for each gene. For each gene with a significant eSTR and eSNP, we analyzed the ability of two models to explain gene expression:

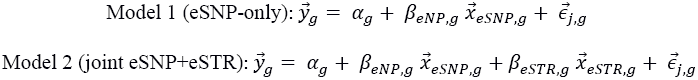

where *α*_*g*_ is the mean expression value for the reference haplotype, 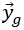 is a vector of expression values for gene *g*, *β*_*eSNP*,*g*_ is the effect of the eSNP on gene *g*, *β*_*eSTR*,*g*_ is the effect of the eSTR on gene *g*, 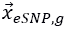 is a vector of genotypes for the best eSNP for gene *g*, 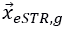 is a vector of genotypes for the best eSTR for gene *g*, and 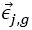 the residual term. A major caveat is that the eSNP dataset has significantly more power to detect associations than the eSTR dataset due to the lower quality of the STR genotype panel (**Supplementary Note**), and this analysis is therefore likely to underestimate the true contribution of STRs to gene expression. We used ANOVA to test whether the joint model performs significantly better than the SNP-only method. We obtained the ANOVA p-value for each gene and used the qvalue package to determine the FDR.

### Conservation analysis

Sequence conservation around STRs was determined using the PhyloP track available from the UCSC Genome Browser. To calculate the significance of the increase in conservation at eSTRs, we compared the mean PhyloP score for each eSTR to that for 1000 random sets of STRs with matched distributions of the distance to the nearest transcription start site. For each STR we determined the mean PhyloP score for a given window size centered on the STR. The p-value given is the percentage of random sets whose mean PhyloP score was greater than the mean of the observed eSTR set.

### Enrichment in histone modification peaks

Chromatin state and histone modification peak annotations generated by the Encode Consortium for GM12878 were downloaded from the UCSC Genome Browser. Because variants involved in regulating gene expression are more likely to fall near genes compared to randomly chosen variants, simple enrichment tests of eSTRs vs. randomly chosen control regions may return strong enrichments simply because of their proximity to genes. To account for this, we randomly shifted the location of eSTRs by a distance drawn from the distribution of distances between the best STR and best SNP for each gene. We repeated this process 1,000 times. For each set of permuted eSTR locations, we generated null distributions by determining the percent of STRs overlapping each annotation. We used these null distributions to calculate empirical p-values for the enrichment of eSTRs in each annotation.

### Predicting effects of STR variation on histone modifications

The WAVE method builds on a kmer-based statistical model to predict the signal of ChIP-seq experiments from a DNA sequence context. Briefly, the model considers that each k-mer has a spatial effect on ChIP-seq read counts in a window of [-M, M-1] bp centered at the start of the k-mer. The read count at a given base is then modeled as the log-linear combination of the effects of all k-mers whose effect ranges cover that base, where k ranges from 1 to 8.

For each eSTR in our dataset, we generated sequences representing each observed allele. We filtered STRs with interruptions in the repeat motif, since the sequence for different allele lengths is ambiguous for these loci. For each mark, we used the model to predict the read count for each allele in a window of ±M bp from the STR boundaries, where M was set to 1,000 for all marks except p300, for which M was set to 200. Previous findings of WAVE showed that these values of M give the best correlation between predicted and real ChIP-seq signals using cross validation. For each alternate allele, we generated a score as the sum of differences in read counts from the reference allele at each position in this window. We regressed the number of repeats for each allele on this score and took the absolute value of the slope for each locus. We repeated the analysis on a set of 2,060 randomly chosen negative control loci and used a Mann-Whitney rank test to compare the magnitudes of slopes between the eSTR and control sets for each mark.

## Acknowledgements

M.G. is supported by the National Defense Science & Engineering Graduate Fellowship. Y.E. holds a Career Award at the Scientific Interface from the Burroughs Wellcome Fund. This study was supported by a gift from Andria and Paul Heafy (Y.E), NIJ grant 2014-DN-BX-K089 (Y.E, T.W), and NIH grants 1U01HG007037 (H.Z), R01MH084703(J.P) and R01HG006399 (A.L.P). We thank Tuuli Lappalainen, Alon Goren, Tatsu Hashimoto, and Dina Zielinksi for useful comments and discussions.

Abundant Contribution of Short Tandem Repeats to gene expression variation in humans Supplementary Material

## 1 Supplementary Methods

### 1.1 Controlling for covariates

We controlled for a number of covariates by regressing them out of the expression dataset. The covariate-corrected expression matrix is given by:

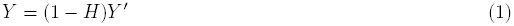

where *Y*′ is an *n × m* matrix of normalized expression values, *Y* is an *n × m* matrix of residualized expression values, *n* is the number of individuals, *m* is the number of genes, *H* = *C*(*C*^*T*^ *C*)^−1^*C*^*T*^ is the hat matrix, and *C* is an *n × c* matrix of *c* covariates. Specifically, the columns of *C* consist of the following sub-matrices:

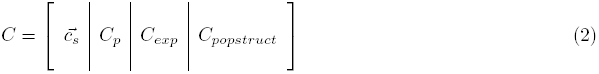

1. **Individual sex**: this is a binary vector, 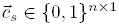, where 0 denotes female and 1 male.
2. **Individual population membership**: this is a binary matrix *C*_*p*_ *∈* {0, 1}^*n×pop-*1^. A “1” in position *C*_*p*_(*i, j*) denotes that individual *i* belongs to population *j*. Specifically, *pop* is equal to to 4 for the association tests with the gEUVADIS RNA-seq data.
3. **Gene expression heterogeneity**: *Y*′ is a matrix that consists of all 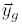 as its column vectors, where 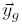 is a vector of expression values for gene *g*. To reduce variation due to experimental differences or other unidentified confounding factors across expression datasets, the top 10 principal components (PCs) corresponding to the top 10 eigenvectors of *Y*′ *Y′^T^* were included as covariates for both the array and RNA-sequencing datasets. *C*_*exp*_ *∈* ℝ^*n×*10^ indicates the matrix of the top 10 PCs.
4. **Population structure**: We first preprocessed the HapMap SNP dataset to include SNPs with MAF *>* 10%. We used Plink [1] for LD-pruning with a pairwise correlation threshold of 0.5, a window size of 50 SNPs, and a step size of 5 SNPs. This left 286,010 SNPs for the RNA-sequencing dataset, which we used to correct for population structure. We used the Tracy-Widom test for population stratification proposed by Patterson, et al. [2] to determine the number of PCs to include as covariates. Let *C*_*popstruct*_ ∈ ℝ^*n×t*^ indicate the matrix of the top *t* PCs removed, where t=5 for the RNA-sequencing dataset.

Residualized expression values were then used as input to the eQTL analysis.

### 1.2 Controlling for gene-level FDR

We controlled for a gene-level false discovery rate (FDR) of 5%, assuming that most genes have at most a single causal eSTR. For each gene, we determined the STR association with the best p-value. This p-value was adjusted using a Bonferonni correction for the number of STRs tested per gene to give a p-value for observing a single eSTR association for each gene. Performing separate permutations for each gene was computationally infeasible, and was found to give similar results to a simple Bonferfonni correction on a subset of genes. We then used this list of adjusted p-values as input to the qvalue package [3] to determine all genes with qval *≤* 5%.

## 2 Supplementary Notes

### 2.1 STR genotype error reduces power to detect eSTRs

We performed simulations to evaluate the effect of lobSTR genotype errors on our power to detect eSTR associations.

We used capillary electrophoresis calls from the Marshfield panel [4] as ground truth genotypes and lobSTR calls for the same markers in our catalog as observed genotypes. We filtered for loci with at least 25 calls for comparison. For each gene, we simulated expression values assuming a single causal STR per gene that explains 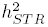 percent of expression variance. We performed the analysis for 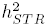 equal to 0.01, 0.05, 0.1, 0.3, and 0.5. Expression values were simulated as follows:

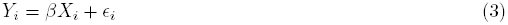

where *Y*_*i*_ is the expression level for individual *i*, *X*_*i*_ is the true STR dosage for individual *i*, 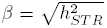 is the effect size of the STR, and 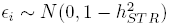 is the residual term for individual *i*.

We performed association analysis regressing 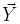 on both 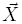 and 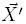, where 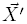 are the observed STR dosages, and tested whether *β* was significantly different than 0 in each case (p*<*0.01). We found that genotype errors limit our power to detect eSTRs (**Supplementary Fig. 1a**) and cause us to underestimate the true variance explained by STRs (**Supplementary Fig. 1b**) but do not introduce spurious eSTR signals.

### 2.2 Comparing expression across array and RNA-Sequencing Datasets

To determine the reproducibility of expression profiling across platforms, we compared gene expression for the 122 individuals profiled by both array and RNA-sequencing. For each platform, we obtained a 122 × 4,627 matrix *Y ^Array^* and *Y*^*RNAseq*^, where 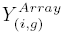 and 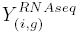 give the expression of gene *g* in individual *i* on the expression array and the RNA sequencing, respectively, before quantile normalization.

We measured the reproducibility of expression profiles inside subjects by calculating the Spearman rank correlation for each pair of row vectors 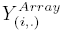 and 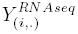 for *i* ∈ {1..122} (**Supplementary Fig. 4a**). The average Spearman correlation was 0.71. A previous study by Maroni et al. [5] measured technical reliability of RNA-seq versus array data with independent datasets. Importantly, they reported an average Spearman correlation of 0.73 for reproducibility of expression profiles inside subjects. This result provides additional support to the technical validity of our expression analysis pipeline.

eQTL replication requires that relative differences between subjects are reproducible across experiments. We compared the order of individuals at each gene as reported by the array and the RNA-sequencing data by measuring the Spearman rank correlation of the column vectors 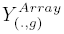 and 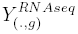 for *g ∈ {*1..4, 627} (**Supplementary Fig. 4b**). The concordance of rank-order of individuals across platforms was moderate (average Spearman rank correlation 0.22), which implies only moderate power to replicate QTLs across the two platforms. Choy et al. performed a similar analysis with biological replicates of LCLs in two expression arrays independent from our study [6]. They also reported Spearman rank correlations of 0.25-0.3 for relative differences of expression between subjects, in agreement with our analysis.

### 2.3 Partitioning heritability on simulated datasets

The best STR can often exhibit high collinearity with other *cis* variants. To rule out the possibility that the LMM could be incorrectly partitioning variance to the STR in the case of tagging another causal variant nearby, we performed simulations in which there was a single causal SNP eQTL per gene. For each gene, we simulated expression values using the following process:

1. Choose the best SNP from the eQTL analysis on real data as the causal variant. Let this eQTL explain *σ*^2^ percent of expression variance.
2. Simulate expression values as *y*_*i*_ = *βx_i_* + *∊*_*i*_ where *y*_*i*_ is the simulated expression value for individual *i*, *x*_*i*_ is the SNP genotype for individual *i*, 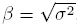, and ∊ ∼ *N*(0, 1 – *σ*^2^).
3. Run the LMM analysis as described above to determine 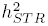 and 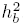.

Notably, this procedure simulates the causal SNP based on the SNP-eQTL analysis, rendering the test more realistic. The simulation was repeated for values of *σ*^2^ equal to 0, 0.01, 0.05, 0.1, 0.2, 0.3, 0.4, and 0.5 for each gene on chromosome 18. We performed this analysis for both the cases of treating the STR as a fixed and a random effect.

We observed that in both models, 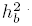 was very close to the simulated value of *σ*^2^, as expected. Importantly, the median value for 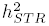 was negative for the fixed effects case and 0 for the random effects case across all simulations. The mean values were close to 0 in most realistic values of SNP-eQTL effects and slightly biased (*<* 0.005) upwards in the case of very strong SNP-eQTLs (**Supplementary Fig. 5**). The median ratio of 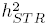 to 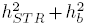 was *<* 0.1% for the fixed effects case and exactly 0 for the random effects case for all simulations. These findings suggest that our LMM analysis reflects an accurate partitioning of variance even in the presence of strong SNP-eQTLs.

To further validate that our estimators of 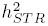 are not inflated, we also ran the fixed effects LMM analysis on random pairs of eSTRs and local bi-allelic mutations from chromosome 2 and gene expression profiles from chromosome 1. This generated a null distribution for 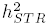 in the case of no association. In this negative control condition, 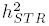 was distributed symmetrically around 0 with mean 7 *×* 10^∔^ and median -0.002, demonstrating that the estimator is unbiased.

### 2.4 Treating STRs as random vs. fixed effects

In our LMM analysis to partition heritability between STRs and other *cis* variants, we treated the best STR for each gene as a fixed effect. We repeated this analysis treating the STR as a random effect to determine whether this choice significantly affects our results. We used a model of the form:

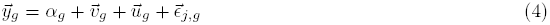

where:

- 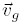 is a length *n* vector of random effects for the best STR
- 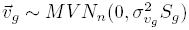 with 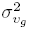 the percent of phenotypic variance explained by the best STR for gene *g*
- *S*_*g*_ is a standardized IBS relatedness matrix constructed using the best STR. It was constructed as:

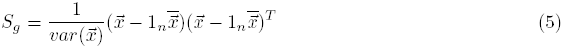

where 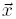 is a length *n* vector consisting of genotypes for the best STR.
- All other variables are as described in the Online Methods.

We used the GCTA program [7] to determine the REML estimates of 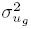 and 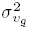. GCTA encountered numerical problems using the 

~~~
--reml-no-constrain
~~~

 option, likely due to the small sample size for each gene and strong correlation between the STR and bi-allelic variance components. Therefore, estimates were constrained to be between 0 and 1 and are biased to be greater than 0.

The overall phenotypic variance-covariance matrix is:

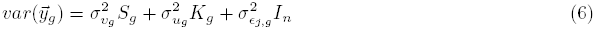

with 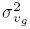 giving the percent of phenotypic variance explained by the best STR 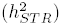 and 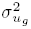 giving the percent explained by other *cis* bi-allelic mutations 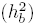.

Estimates of the variance explained by STRs and by *cis* bi-allelic mutations using this model are consistent with those obtained by treating STRs as a fixed effect (**Table 1** and **Supplementary Table 5**). Because the random effects estimates are constrained to be between 0 and 1, the random effects model tended to partition variance all to a single variance component, but overall distributions of 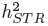 and 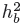 were similar to the fixed effects case (**Fig. 3a,b** and **Supplementary Fig. 6**).

## 3 Supplementary Figures

**3.1 Supplementary Figure 1:**
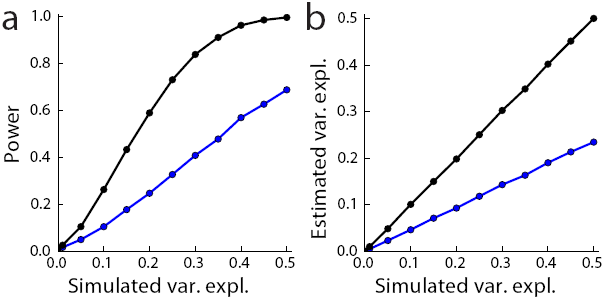
STR genotyping errors reduce power to detect eSTR associations. **a**. Power to detect associations and **b**. estimated variance explained for different simulated values of variance explained by the STR. (black: observed capillary electrophoresis genotypes, blue: lobSTR genotypes).

**3.2 Supplementary Figure 2:**
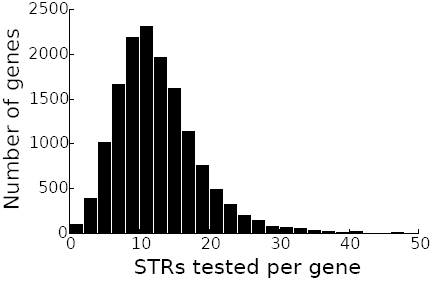
Number of sTRS tested per gene. Histogram gives the number of STRs within 100kb of each gene that passed quality filters and were included in the eSTR analysis.

**3.3 Supplementary Figure 3:**
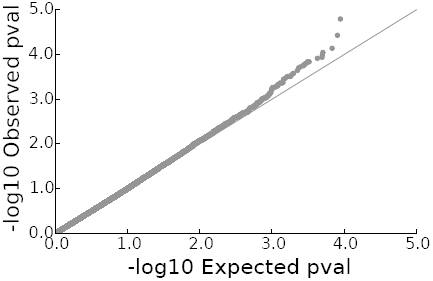
Unlinked controls follow the null. QQ plot of association tests between random unlinked STRs and genes.

**3.4 Supplementary Figure 4:**
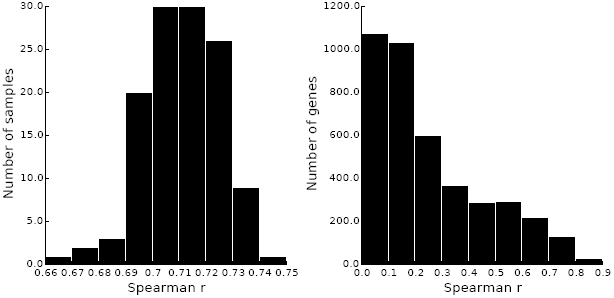
Expression values are moderately reproducible across platforms. **a**. Distribution of Spearman rank correlation coefficients between gene expression profiles of individuals measured on microarray vs. RNA-sequencing platforms. **b**. Distribution of Spearman rank correlation coefficients between the order of individuals ranked by expression levels across transcripts measured using microarray vs. RNA-sequencing platforms.

**3.5 Supplementary Figure 5:**
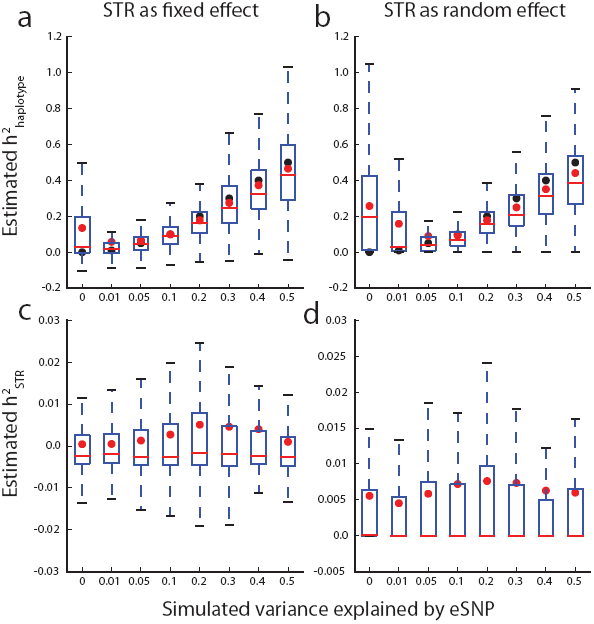
Variance partitioning simulations. Plots show variance partitioning results from simulations in which each gene has a single causal eSNP. (**a**&**b**) The distributions of 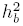 The distributions of 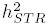 (**a**&**c**) *b ST R* The LMM simulations with STRs as fixed effects (**b**&**d**) The LMM simulations with STRs as random effects (**a**-**d**) Black points denote the true value of the variance explained by the causal SNP. Red dots denote the average value of the estimator. Red bars denote the median value of the estimator. The figure shows that the median values of the best STRs are insensitive to the presence of a strong SNP eQTL.

**3.6 Supplementary Figure 6:**
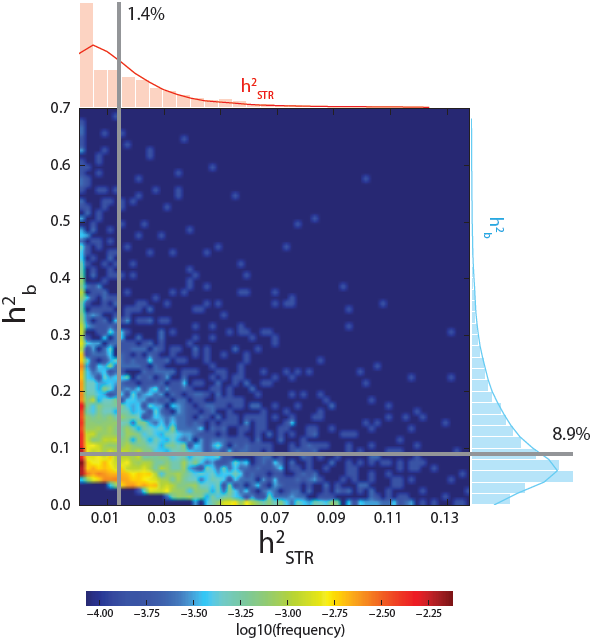
Partitioning variance when treating the STR as a random effect. The heatmap shows the distribution 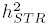 and 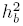 for each gene. Dashed gray lines give the medians of each distribution.

**3.7 Supplementary Figure 7:**
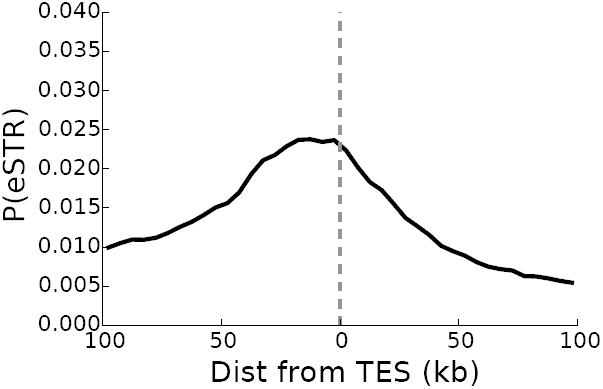
Enrichment of eSTRs at transcription end sites eSTRs are enriched near the transcription end site (TES). For each distance bin around the TES, the plot shows the percentage of STRs in that bin that were called as significant eSTRs.

## 4 Supplementary Tables

**4.1 SUPPLEMENTARY TABLE 1: SIGNIFICANT ESTRS**

See file Gymrek etal signifcant estrs.tab.

**4.2 Supplementary Table 2:** Distribution of motif lengths in eSTRs vs. all STRs. Distribution of motif lengths in all unique eSTR loci vs. all unique STR loci included in the analysis after applying quality filters.

| Period | Num. eSTRs | % eSTRs | Num. all STRs | % all STRs | Enrichment | Pval |
| --- | --- | --- | --- | --- | --- | --- |
| 2 | 951 | 50.2% | 50,184 | 62.0% | 0.81 | 1.0 |
| 3 | 223 | 11.8% | 7,369 | 9.1% | 1.29 | $4.8 \times 10^{-5}$ |
| 4 | 516 | 27.2% | 17,938 | 22.2% | 1.23 | $8.2 \times 10^{-8}$ |
| 5 | 166 | 8.8% | 4,466 | 5.5% | 1.59 | $3.9 \times 10^{-9}$ |
| 6 | 39 | 2.1% | 1,023 | 1.3% | 1.63 | $2.4 \times 10^{-3}$ |

**4.3 Supplementary Table 3:**
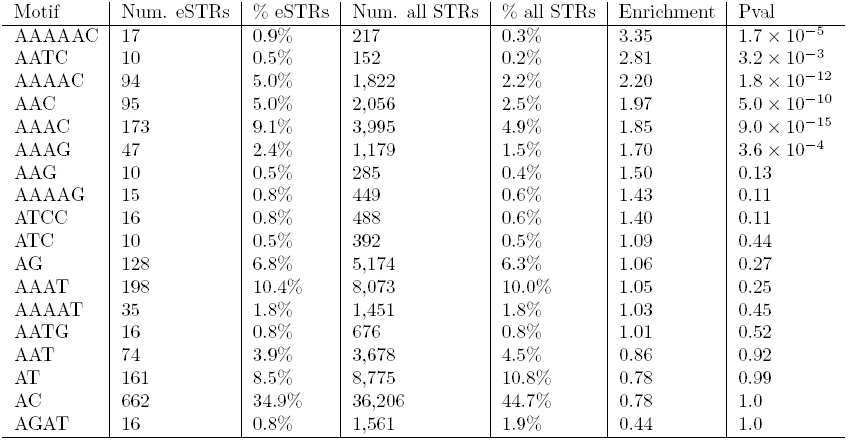
Distribution of motifs in eSTRs vs. all STRs. Distribution of motifs in all unique eSTR loci vs. all unique STR loci included in the analysis after applying quality filters. Only motifs for which there were at least 10 eSTRs are shown. Motifs were converted to canonical format as described in [8].

| Motif | Num. eSTRs | % eSTRs | Num. all STRs | % all STRs | Enrichment | Pval |
| --- | --- | --- | --- | --- | --- | --- |
| AAAAAC | 17 | 0.9% | 217 | 0.3% | 3.35 | $1.7 \times 10^{-5}$ |
| AATC | 10 | 0.5% | 152 | 0.2% | 2.81 | $3.2 \times 10^{-3}$ |
| AAAAC | 94 | 5.0% | 1,822 | 2.2% | 2.20 | $1.8 \times 10^{-12}$ |
| AAC | 95 | 5.0% | 2,056 | 2.5% | 1.97 | $5.0 \times 10^{-10}$ |
| AAAC | 173 | 9.1% | 3,995 | 4.9% | 1.85 | $9.0 \times 10^{-15}$ |
| AAAG | 47 | 2.4% | 1,179 | 1.5% | 1.70 | $3.6 \times 10^{-4}$ |
| AAG | 10 | 0.5% | 285 | 0.4% | 1.50 | 0.13 |
| AAAAG | 15 | 0.8% | 449 | 0.6% | 1.43 | 0.11 |
| ATCC | 16 | 0.8% | 488 | 0.6% | 1.40 | 0.11 |
| ATC | 10 | 0.5% | 392 | 0.5% | 1.09 | 0.44 |
| AG | 128 | 6.8% | 5,174 | 6.3% | 1.06 | 0.27 |
| AAAT | 198 | 10.4% | 8,073 | 10.0% | 1.05 | 0.25 |
| AAAAT | 35 | 1.8% | 1,451 | 1.8% | 1.03 | 0.45 |
| AATG | 16 | 0.8% | 676 | 0.8% | 1.01 | 0.52 |
| AAT | 74 | 3.9% | 3,678 | 4.5% | 0.86 | 0.92 |
| AT | 161 | 8.5% | 8,775 | 10.8% | 0.78 | 0.99 |
| AC | 662 | 34.9% | 36,206 | 44.7% | 0.78 | 1.0 |
| AGAT | 16 | 0.8% | 1,561 | 1.9% | 0.44 | 1.0 |

**4.4 Supplementary Table 4:** Distribution of genomic locations of eSTRs vs. all STRs. Annotations were compiled using Ensembl version 71. “Exon” refers to both coding and non-coding exons and untranslated regions. “Neargene” refers to regions within of a gene. “Intergenic” refers to STRs not falling into any other annotation. Note some STRs may overlap multiple annotations.

| Annotation | Num. eSTRs | % eSTRs | Num. all STRs | % all STRs | Enrichment | Pval |
| --- | --- | --- | --- | --- | --- | --- |
| Coding | 13 | 0.7% | 157 | 0.2% | 3.54 | $9.1 \times 10^{-5}$ |
| 5' UTR | 51 | 2.7% | 897 | 1.1% | 2.43 | $1.0 \times 10^{-8}$ |
| Exon | 127 | 6.7% | 2,452 | 3.0% | 2.21 | $1.5 \times 10^{-16}$ |
| 3' UTR | 77 | 4.1% | 1,569 | 1.9% | 2.10 | $1.7 \times 10^{-9}$ |
| Neargene (5') | 335 | 17.7% | 7,357 | 9.1% | 1.95 | $1.5 \times 10^{-32}$ |
| Neargene (3') | 326 | 17.2% | 7,399 | 9.1% | 1.88 | $4.5 \times 10^{-29}$ |
| Intron | 1,314 | 69.3% | 52,326 | 64.6% | 1.07 | $6.1 \times 10^{-6}$ |
| Intergenic | 395 | 20.8% | 23,373 | 28.9% | 0.72 | 1.00 |

**4.5 Supplementary Table 5:** Heritability of gene expression explained by STRs vs. SNPs in each LMM. Values show the median and 95% confidence interval of the median across all eSTR-containing genes and genes with moderate *cis* heritability (*≥*5%). 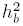 denotes the variance explained by all common *cis* bi-allelic markers, 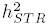 denotes the variance explained by the best STR for each gene, and 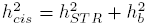.

| | $h_b^2$ | $h_{STR}^2$ | $h_{STR}^2/h_{cis}^2$ |
| --- | --- | --- | --- |
| eSTR genes - (STR fixed) | 0.1203 (0.1139-0.1259) | 0.0180 (0.0166-0.0199) | 0.1230 (0.1106-0.1420) |
| eSTR genes - (STR random) | 0.1229 (0.1159-0.1295) | 0.0200 (0.0178-0.0216) | 0.1288 (0.1179-0.1451) |
| Moderate *cis* $h_{cis}^2$ (STR fixed) | 0.0910 (0.0884-0.0938) | 0.0145 (0.0137-0.0151) | 0.1283 (0.1222-0.1346) |
| Moderate *cis* $h_{cis}^2$ (STR random) | 0.0892 (0.0865-0.0918) | 0.0143 (0.0137-0.0149) | 0.1245 (0.1184-0.1309) |

